# Enteroendocrine Cells Regulate Intestinal Barrier Permeability

**DOI:** 10.1101/2025.03.07.642036

**Authors:** Jennifer G. Nwako, Sparsh D. Patel, Taevon J. Roach, Saanvi R. Gupte, Samara G. Williams, Anne Marie Riedman, Heather A. McCauley

**Author notes:** Correspondence to: Heather A. McCauley, Ph.D., ^1^Department of Cell Biology and Physiology, University of North Carolina at Chapel Hill School of Medicine, 111 Mason Farm Road, Molecular Biology Research Building 5341C, Chapel Hill, NC 27599.

## Abstract

The intestinal epithelial barrier is essential for nutrient absorption and protection against ingested pathogens and foreign substances. Barrier integrity is maintained by tight junctions which are sensitive to inflammatory signals, thus creating a feed-forward loop with an increasingly permeable barrier that further drives inflammation and is the hallmark of inflammatory bowel disease. There are currently no therapeutic strategies to improve the intestinal epithelial barrier. We hypothesized that enteroendocrine cells may play an unappreciated role in maintaining barrier integrity. To test this hypothesis, we seeded human intestinal enteroids with genetic loss of enteroendocrine cells on Transwell filters and evaluated transepithelial electrical resistance, paracellular permeability, and the localization and abundance of junctional proteins. We found that enteroendocrine cells were required to maintain a healthy barrier in crypt-like “stem” and villus-like differentiated cultures. Additionally, exogenous supplementation of enteroendocrine-deficient cultures with the hormones peptide tyrosine tyrosine (PYY) and the somatostatin analog octreotide was sufficient to rescue many aspects of this barrier defect both at baseline and in the presence of the inflammatory cytokine tumor necrosis factor (TNF). Surprisingly, these improvements in barrier function occurred largely independently of changes in protein abundance of junctional proteins zona-occludens 1, occludin, and claudin-2. These findings support a novel role for enteroendocrine cells in augmenting epithelial barrier function in the presence of inflammatory stimuli and present an opportunity for developing therapies to improve the intestinal barrier.

**NEW & NOTEWORTHY:** There are no therapies that directly improve the permeability of the intestinal epithelial barrier. This work uses a human intestinal epithelial model system to demonstrate that sensory enteroendocrine cells are necessary for healthy barrier function and that two of their secreted products, peptide YY and somatostatin, are sufficient to improve barrier function at homeostasis and in the presence of inflammatory cytokines. This could provide novel treatments for strengthening the epithelial barrier in human gastrointestinal disease.

## INTRODUCTION

The intestinal epithelial barrier performs two essential functions: absorption of nutrients and defense against pathogens and bacteria. Intestinal inflammation stemming from a disrupted barrier is the hallmark of inflammatory bowel disease (IBD) and is associated with increased expression of the pro-inflammatory cytokine tumor necrosis factor (TNF) (1). TNF further damages the epithelial barrier by disrupting tight junctional permeability (2-4). This damage results from the degradation of tight junction proteins that support a healthy barrier such as ZO-1 (zona occludens 1), occludin, and junctional adhesion molecules (JAMs). Additionally, junctional proteins that promote a more permeable barrier, such as the pore-forming claudin-2, are upregulated in IBD (5, 6). This creates a vicious cycle which worsens patient outcomes (1).

Anti-TNF biologic therapies currently offer the best clinical outcome for people with IBD. However, not all patients respond to this type of therapy, and there are currently no therapies aimed at bolstering the epithelial barrier. Strengthening the junctional proteins that comprise the epithelial barrier could help naturally resolve excessive pro-inflammatory cytokine release by limiting the intestinal permeability to foreign substances.

Here, we investigate whether a specialized intestinal epithelial cell, the enteroendocrine cell (EEC), participates in intestinal barrier permeability. EECs are sensory cells that secrete hormones, metabolites, and other small molecules in response to cues such as nutrients and microbes. While classically known for their systemic digestive and metabolic actions, the roles of EECs within the intestine are only beginning to be understood, largely due to the lack of tractable model systems(7). The localization of receptors for EEC hormones gives some clues as to their potential functions, with receptors for many hormones, including peptide tyrosine tyrosine (PYY) and somatostatin, found on intestinal epithelial cells(8).

We previously developed a novel model system to experimentally test the role of EECs in intestinal physiology and function by generating human intestinal organoids (HIOs) from pluripotent stem cells with a null mutation in *NEUROG3*, the transcription factor required for EEC differentiation(9). After maturation by xenograft into mice, we isolated crypts from wild-type and EEC-deficient HIOs for enteroid culture. The resulting enteroids differentiate into expected intestinal cell populations, including EECs in wild-type enteroids. As anticipated, EEC-deficient enteroids fail to generate EECs or their products, like PYY and somatostatin. We previously used this model to demonstrate that EECs are necessary to regulate ion-coupled nutrient absorption(9) and crypt cell metabolism(10) in the small intestine.

In this study, we used EEC-deficient enteroids to test the hypothesis that EECs would also be important regulators of barrier permeability, the other essential function of the intestinal epithelium. We found that loss of all EECs resulted in impaired barrier function, whereas EEC-derived hormones PYY and somatostatin were each sufficient to improve barrier function in homeostatic conditions and in the context of TNF-mediated barrier dysfunction. These findings elucidate a novel role for EECs in intestinal physiology and pathophysiology and may form the basis of future therapies aimed at strengthening the intestinal epithelial barrier.

## MATERIALS & METHODS

### Cell Culture

Expansion media (EM) and differentiation media (DM) were prepared as previously described(11). Wild-type and EEC-deficient (NEUROG3-/-) human intestinal enteroids were cultured and maintained as previously described(9). Cells were dissociated and plated in monolayer on collagen patties and on Transwell filters (Fisher #07-200-154) as previously described (11, 12). After reaching confluency in EM (approximately 3 days), cells were switched to DM for an additional 7 days.

### OCT, PYY, and TNF Treatment *in vitro*

Cells were seeded and maintained on Transwell filters until the cells formed a confluent monolayer as measured by a plateau in TEER values. Human recombinant TNF, 150 ng/Ml (PeproTech #300-01A-100UG), octreotide, 10 μM (OCT; Sigma-Aldrich #01014-1MG) or human PYY, 1 μM (Phoenix Pharmaceuticals #059-07) were added to the basal Transwell chamber in the appropriate media and incubated for 24 hours at 37°C, 5% CO_2_. Cell culture medium served as vehicle control.

### Trans-epithelial Electrical Resistance (TEER)

TEER measurements were performed using an EVOM Manual (EVM-MT-03-01) according to manufacturer’s instructions. Empty Transwell filters coated as described and containing only media were used as blanks. Resistance values (Ω) were recorded after subtracting the resistance contribution of the blank filter. A minimum of 5 composite wells in each group was recorded to reduce experimental errors.

### Barrier Permeability

Cells on Transwell filters were incubated on the apical side only in HBSS containing 1M HEPES buffer at pH 7.4 and 0.1 mg/ml Lucifer Yellow (LY) CH dipotassium salt (Molecular Probes #L1177). Buffer without LY was added to the basolateral side. Cells were incubated for 2 hours at 37°C prior to collecting media from the basal compartment for analysis. The fluorescence signal (excitation at 485 nm and emission at 538 nm) was measured using a plate reader (Clariostar Plus, BMG Labtech) and LY percent permeability was calculated based on fluorescence intensity. Protocol adjusted from Sigma Aldrich (Technical Bulletin MTOX1000P24).

### Western Blot

Protein was isolated from cell pellets of enteroid monolayers and subjected to SDS-PAGE using standard procedures (Bio-rad Bulletin #6376). After transfer, the nitrocellulose membrane was blocked, stained, and imaged using LI-COR reagents and according to their protocol. The primary antibodies include the following: Claudin 2 (1:200; Cell Signaling; #48120S), Zona occludens 1 (1:500; Proteintech Group, Inc; #10019107), Occludin (1:250; #71-1500; Invitrogen) GAPDH (1:2500; #GTX627408; GeneTex). LI-COR secondary antibodies at 1:15,000 dilution include Goat anti-rabbit IgG (#926-68071; #926-32211) and Goat anti-mouse IgG (#926-32210, #926-68070). Western Blots were quantified in Image J software with GAPDH serving as loading control. Experiments were repeated in triplicate.

### Immunofluorescence

Human enteroid monolayers on Transwell filters were fixed and stained as previously described(13). The primary antibodies used were: Claudin 2 (1:100; Invitrogen; #32-5600), Zona occludens 1 (1:100; Invitrogen; #33-9100), Occludin (1:100; #33-1500; Invitrogen). All secondary antibodies were conjugated to Alexa Fluor 488 or 647 (Invitrogen, #A-21202, #A-31573) and used at 1:1000 dilution. The Transwell membrane was then removed from the plastic insert and mounted in Fluoromount with DAPI (Fisher Scientific #501128966) overnight at room temperature prior to imaging.

### Confocal Microscopy

Confocal images were acquired using a Zeiss LSM 900 running Zen 2019 imaging software, blue edition (Zeiss). Images were captured with a 63x oil immersion objective. For qualitative display, image settings were optimally adjusted, and the same settings were used to image across samples in Image J. We quantified the fluorescence intensity of the maximum intensity projection of z-stacked images using a macro in Image J after subtracting background signal. Experiments were conducted in triplicate.

### Statistical Analysis

All experiments were conducted with a minimum of n=5. Results are expressed as the mean ± standard error of the mean. GraphPad Prism 10 software was used for statistical analysis. P-values were calculated using one-way ANOVA with Tukey’s multiple comparisons test or unpaired two-tailed Student’s t-test as appropriate. P<0.05 was considered to indicate a statistically significant difference, and P-values are reported in each figure.

## RESULTS

EECs are found throughout the crypt-villus axis in vivo and we hypothesized they may have different effects on target cells based on location and function. Tight junctions are looser in the crypt and strengthen as cells differentiate in the villus; this is recapitulated in enteroid culture when comparing TEER values between undifferentiated, “stem”-like cultures and differentiated cultures(14). Enteroids were plated in monolayer culture on a Transwell filter and allowed to grow to confluency in high-Wnt media, at which point we either performed analysis in stem-like conditions or switched the cultures to low-Wnt media to promote differentiation over the following 7 days (Figure 1A). We measured and quantified barrier function in two ways: by TEER and by paracellular permeability to the small molecule Lucifer Yellow (LY, 457 Da). We found a small but significant decrease in TEER in stem-like EEC-deficient enteroids compared to wild-type control (Figure 1B) and equivalent permeability to LY between stem-like wild-type and EEC-deficient enteroid cultures (Figure 1C). In EEC-deficient differentiated cultures, we found a dramatic reduction in TEER (Figure 1D) and increased permeability to LY (Figure 1E) compared to wild-type, suggesting that EECs and/or their secreted products were important in maintaining a tight intestinal epithelial barrier.

**Figure 1.**
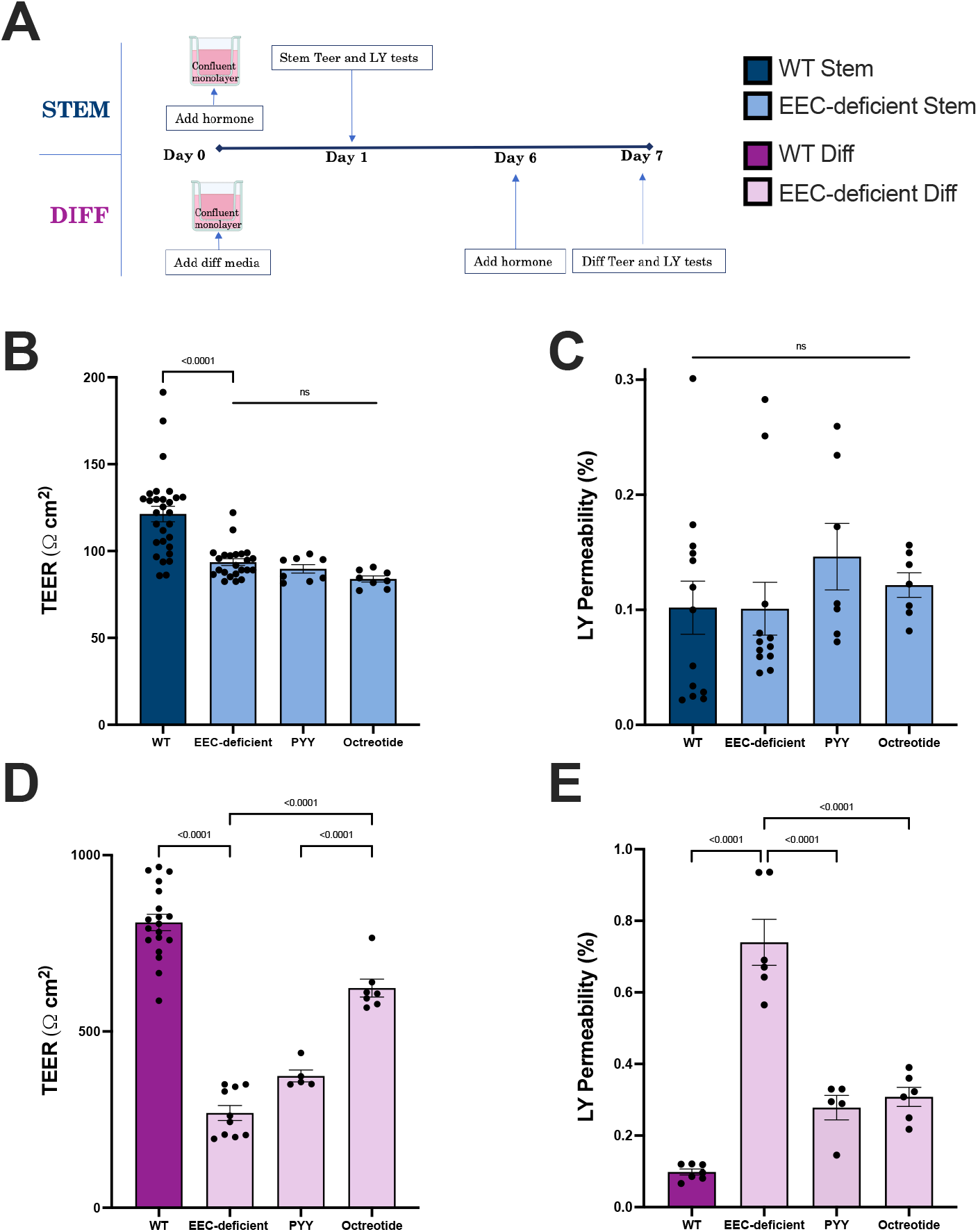
EECs regulate barrier function in the intestinal epithelium. (A) Enteroids were seeded onto semi-permeable Transwell filters to generate 2D cultures. When monolayers were confluent, they were randomly allocated for analysis of stem or differentiated populations. Hormones were added 24 hours prior to barrier function testing. (B, D) TEER and (C, E) permeability to Lucifer Yellow were measured in WT and EEC-deficient enteroid Transwell monolayers in stem (B,C) and differentiated (D,E) conditions. Data reported as mean ± SEM. P-values calculated by one-way ANOVA with Tukey’s multiple comparison’s test. ns, not significant. *n* = 5-30 wells/group.

We next exploited the absence of all EECs in EEC-deficient enteroids to systematically test the ability of individual EEC hormones to impact barrier function. Octreotide is a synthetic analog to somatostatin which is clinically approved for the treatment of diarrhea and has been shown to regulate the expression of junctional proteins (15) and strengthen the intestinal epithelial barrier in mice with experimental colitis (16) and the Caco2 cell line (17, 18). Alongside octreotide, we hypothesized that PYY might improve TEER and permeability to LY in human EEC-deficient enteroid monolayers. Like somatostatin, PYY is an inhibitory enteroendocrine peptide that we have previously demonstrated regulates intestinal epithelial functions in mouse and human small intestine (9). We exposed EEC-deficient enteroid monolayers to octreotide and PYY in the basal chamber for 24 hours preceding analysis (Figure 1A). Both octreotide and PYY robustly improved TEER (Figure 1D) and permeability to LY (Figure 1E) in EEC-deficient differentiated cultures, although neither hormone impacted barrier function in EEC-deficient stem-like cultures (Figure 1B, C).

As inflammatory cytokines like TNF are well-known to disrupt the intestinal epithelial barrier (2, 4), we next evaluated whether EECs protected against TNF-mediated barrier dysfunction. While lower doses of TNF were sufficient to elicit a response in differentiated cultures (data not shown), both wild-type and EEC-deficient stem-like cultures required a higher (150 ng/mL) dose of TNF to significantly alter TEER and permeability. For consistency, we proceeded to treat all cultures with the same concentrations of TNF, PYY, and octreotide (Figure 2). Treatment of wild-type and EEC-deficient stem and differentiated cultures with TNF for 24 hours in the basal chamber significantly disrupted TEER (Figure 2 A, C). TNF caused increased permeability to LY in differentiated wild-type and EEC-deficient cultures (Figure 2D), but wild-type stem-like cultures were relatively protected against increased permeability even at a high dose whereas EEC-deficient stem-like cultures were more sensitive to TNF (Figure 2B). When we treated EEC-deficient cultures concurrently with octreotide or PYY alongside TNF, we found that these two EEC-derived hormones impacted inflammation-mediated barrier function in disparate ways. Both octreotide and PYY improved permeability to LY in stem cultures (Figure 2B) despite not affecting TEER (Figure 2A). Interestingly, in TNF-treated EEC-deficient differentiated cultures, octreotide improved TEER but not permeability to LY, whereas PYY treatment rescued both TEER and permeability to near wild-type levels (Figure 2C, D). This suggested that somatostatin and PYY may impact tight junctions by different mechanisms.

**Figure 2.**
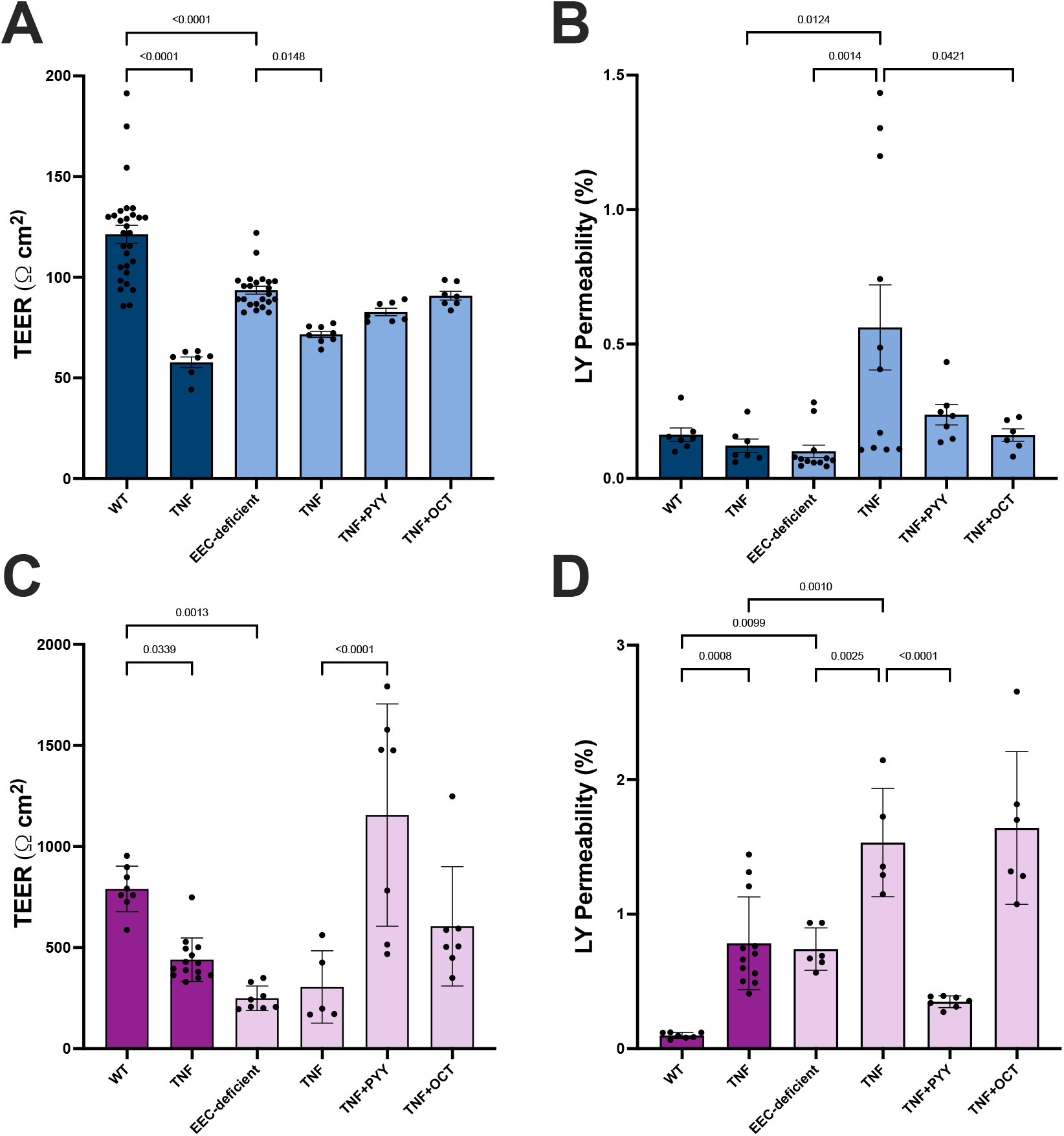
EECs improve TNF-mediated barrier dysfunction. Stem and differentiated EEC-deficient enteroid monolayers were treated with octreotide (10 μM), and/or PYY (1 μM) concurrently with 150 ng/mL TNF for 24 hours prior to measuring (A, C) TEER and (B, D) permeability to Lucifer Yellow. Data reported as mean ± SEM. P-values calculated by one-way ANOVA with Tukey’s multiple comparison’s test. ns, not significant. *n* = 5-30 wells/group.

We next sought to visualize and quantify the impact of octreotide and PYY on tight junctional proteins with and without the inflammatory cytokine TNF. We observed immunofluorescence expression of ZO-1 and claudin-2 in wild-type and EEC-deficient stem-like cultures regardless of the addition of TNF (Figure 3A). Qualitatively, we noted punctate staining of claudin-2 in wild-type stem cultures, which appeared thickened and more evenly distributed throughout the cell membranes of EEC-deficient stem cultures (Figure 3A). In contrast, the smooth distribution of ZO-1 along wild-type cell membranes appeared weakened in EEC-deficient stem cultures (Figure 3A). These qualitative assessments were supported by a significant decrease in fluorescence intensity and total protein of ZO-1 (Figure 3A, B) and total protein of occludin (Figure 3C) in EEC-deficient stem cultures compared to wild-type, with a trend toward an increase in the pore-forming claudin-2 (Figure 3A, D). Exogenous addition of PYY or octreotide restored ZO-1 and occludin to wild-type levels in EEC-deficient stem cultures (Figure 3B, C) but did not affect expression of claudin-2 (Figure D). Interestingly, while TNF treatment functionally disrupted TEER and permeability (Figure 2A, B), and qualitatively altered the sharp membrane expression of ZO-1 and claudin-2 (Figure 3A), these effects were not reliant on changes in total expression of tight junctional proteins and were not impacted by exogenous hormone treatment in EEC-deficient stem cultures (Figure 3B-D).

**Figure 3.**
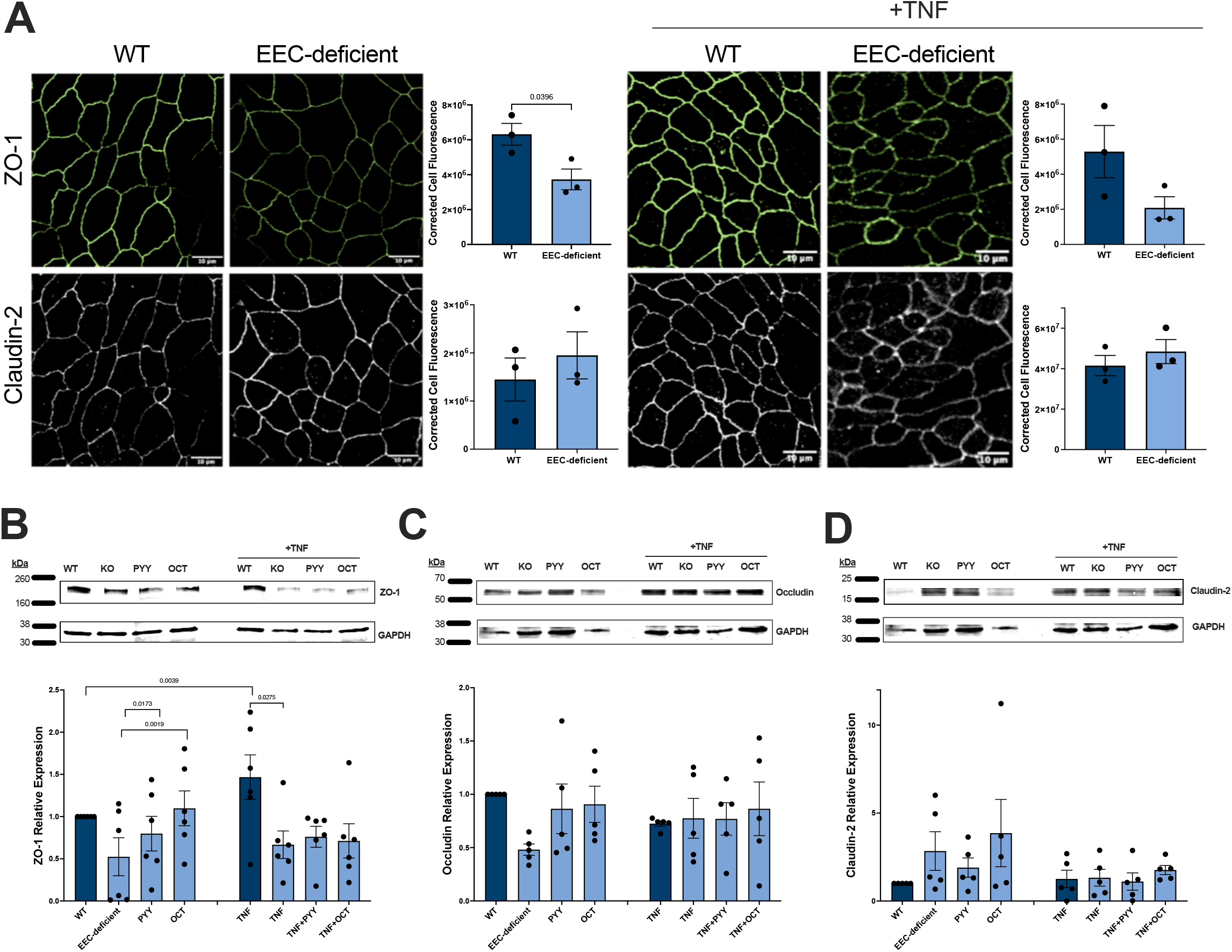
Tight junction proteins are altered in crypt-like cells upon the loss of EECs. (A) Immunofluorescence staining and quantification of corrected cell total fluorescence intensity for ZO-1 and Claudin-2 in enteroid monolayers grown in stem conditions with and without treatment with TNF. Scale bars = 10 μm. (B-D) EEC-deficient enteroid monolayers grown in stem conditions were treated with octreotide (10 μM), and/or PYY (1 μM) concurrently with 150 ng/mL TNF for 24 hours prior to quantification of (B) ZO-1 (225 kDa), (C) occludin (55 kDa), and (D) claudin-2 (22 kDa) protein abundance by Western blot. Data reported as mean ± SEM. P-values calculated by unpaired *t*-test. *n =* 3 immunofluorescence images per condition; *n* = 5-6 immunoblots per condition.

Similarly, we observed a qualitative and quantitative decrease of ZO-1 and occludin in EEC-deficient differentiated cultures compared to wild-type (Figure 4A-C). Consistent with the preferential crypt localization of claudin-2(19), we did not detect claudin-2 expression in our differentiated enteroid cultures (not shown). Treatment with TNF resulted in thickened and interrupted membrane staining in both wild-type and EEC-deficient differentiated cultures (Figure 4A), although these qualitative changes did not correlate with statistically significant changes in protein levels (Figure 4A-C). Similar to stem cultures, neither PYY nor octreotide increased ZO-1 or occludin protein levels in TNF-treated differentiated EEC-deficient cultures (Figure 4B, C).

**Figure 4.**
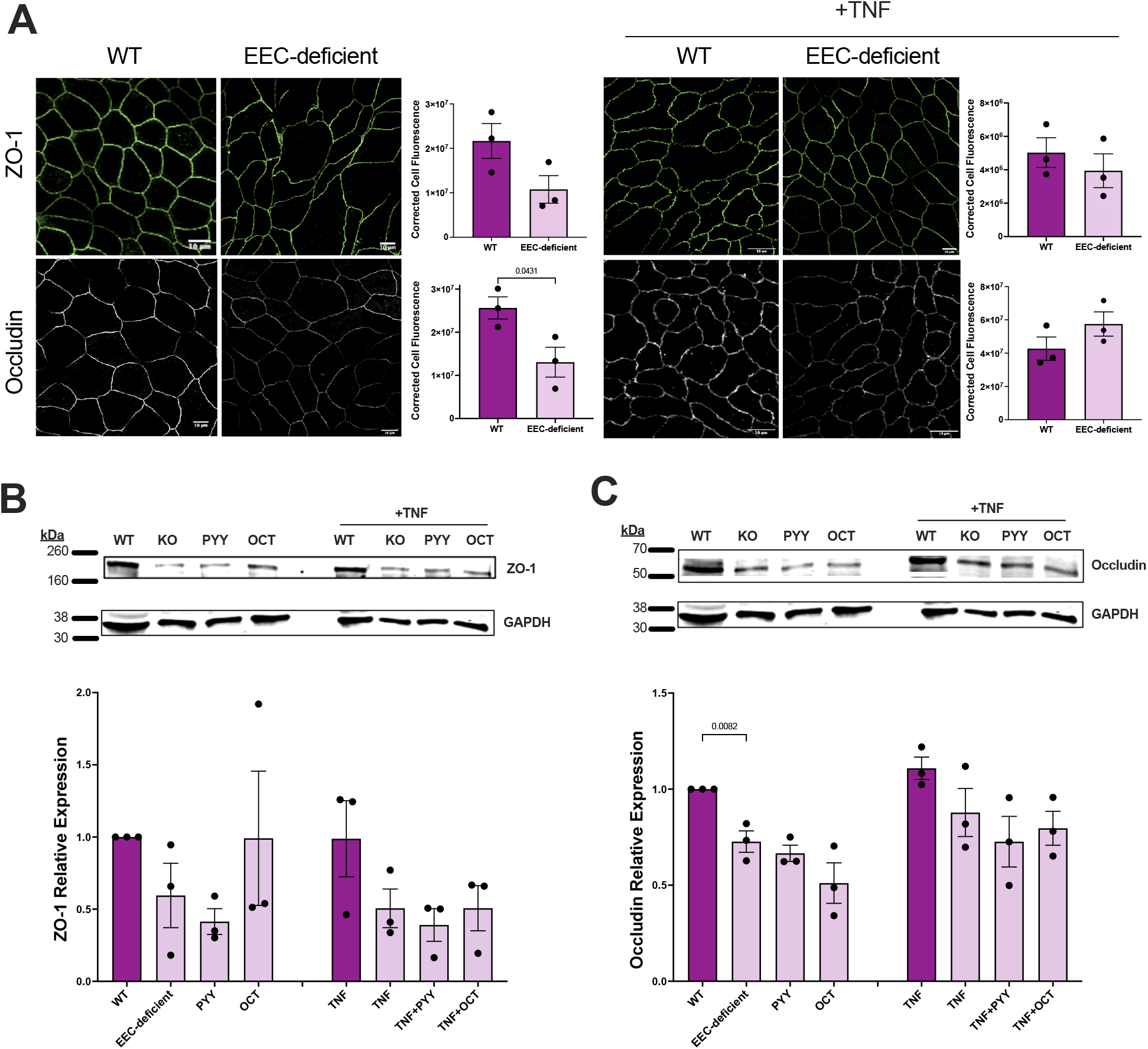
Tight junction proteins are altered in villus-like cells upon the loss of EECs. (A) Immunofluorescence staining and quantification of corrected cell total fluorescence intensity for ZO-1 and occludin in enteroid monolayers after 7 days of differentiation with and without treatment with TNF during the final 24 hours. Scale bars = 10 μm. (B-D) Differentiated EEC-deficient enteroid monolayers were treated with octreotide (10 μM), and/or PYY (1 μM) concurrently with 150 ng/mL TNF for 24 hours prior to quantification of (B) ZO-1 (225 kDa), and (C) occludin (55 kDa) protein abundance by Western blot. Data reported as mean ± SEM. P-values calculated by unpaired *t*-test. *n =* 3 immunofluorescence images per condition; *n* = 3 immunoblots per condition.

Taken together, our results demonstrate a novel role for EECs and their secreted products in maintaining the integrity of the intestinal epithelial barrier in a manner that is largely independent of quantitative changes in protein expression of tight junction family members.

## DISCUSSION

In this study, we demonstrated that EECs support human intestinal epithelial barrier function and that two of their secreted products, PYY and somatostatin, were sufficient to improve barrier function at homeostasis and in the presence of high levels of the pro-inflammatory cytokine TNF. To our knowledge, this is the first report demonstrating the role of PYY, a classical satiety hormone, in strengthening the intestinal barrier.

We investigated how EECs impacted the epithelial barrier in crypt-like “stem” and villus-like differentiated cultures. In EEC-deficient cultures, we found a slight but significant reduction in TEER in stem-like cultures with a concomitant decrease of ZO-1 protein expression and increase in the pore-forming protein claudin-2 compared to wild-type. In differentiated cultures, loss of EECs resulted in a “leaky” barrier with decreased TEER and increased permeability compared to wild-type cultures. We observed decreased ZO-1 and occludin expression in EEC-deficient cultures that, surprisingly, was not further exacerbated by TNF treatment. Interestingly, while PYY and octreotide were both able to improve TEER in EEC-deficient TNF-treated differentiated cultures, only PYY was able to improve permeability, and neither hormone impacted protein levels of ZO-1 or occludin. This, coupled with the qualitative changes in ZO-1, occludin, and claudin-2 expression and localization we observed by immunofluorescence, suggests that PYY and somatostatin are sufficient to mitigate epithelial damage through unexplored mechanisms independent of junctional protein abundance. Future experiments will investigate the role of EECs in tight junction ultrastructure, endocytosis protein recycling, and junctional complex interactions with additional stabilizing proteins, like the myosin light chain, and the lipid membrane. It is also possible that individual hormones might impact these processes differently in the crypt versus the villus.

As neither wild-type nor EEC-deficient enteroids form EECs in the media conditions that support the stem-like cultures, we had expected roughly equivalent characteristics between lines. Baseline differences in TEER and junctional proteins raise the possibility that wild-type intestinal stem cells retain some epigenetic memory of their previous exposure to EEC hormones *in vivo*. Our data suggest that EEC-deficient stem-like cultures are primed for increased permeability that is exacerbated upon exposure to inflammatory cytokines.

Here, we provide proof-of-concept that EECs may be targeted for improving the integrity and function of the intestinal epithelial barrier. EECs and their secreted hormones are often dysregulated in GI diseases; for example, colonic biopsies from patients with severe Crohn’s disease and ulcerative colitis revealed loss of PYY- and somatostatin-positive cells (20-22). While it is unclear whether alterations in EECs drive disease or are merely an effect of a damaged epithelium, our study demonstrates a theoretical basis for supplementing PYY and/or somatostatin in these patients in an effort to reduce barrier permeability and improve disease outcomes.

## ACKNOWLEDGMENTS

We thank the UNC Center for Gastrointestinal Biology and Disease (CGIBD), especially the Advanced Analytics Core, for assistance with this project. The Graphical Abstract was created with BioRender.com.

## AUTHOR CONTRIBUTIONS

J.G.N. designed research, performed experiments, analyzed data, interpreted results of experiments, prepared figures, drafted manuscript, edited and revised manuscript, and approved final version of manuscript. S.D.P., T.J.R., S.R.G., S.G.W., and A.M.R. performed experiments, edited and revised manuscript, and approved final version of manuscript. H.A.M. conceived and designed research, performed experiments, analyzed data, interpreted results of experiments, prepared figures, drafted manuscript, edited and revised manuscript, and approved final version of manuscript.

---

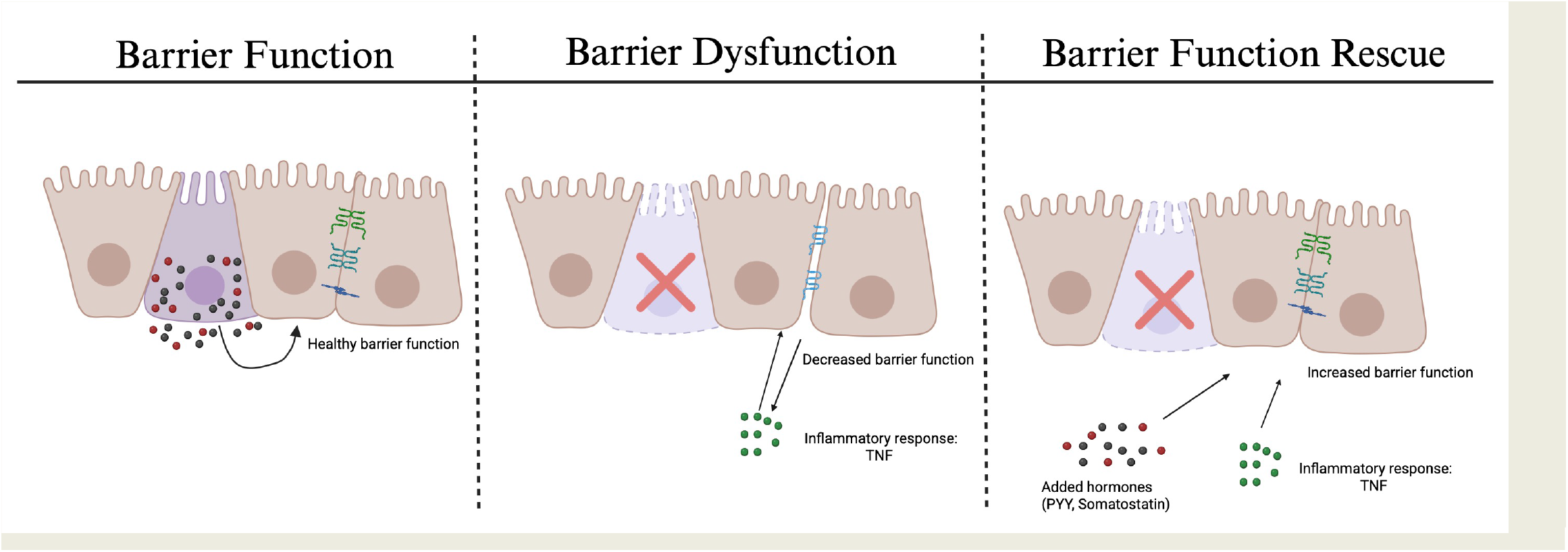

